# Differential effects of cerebellar degeneration on feedforward versus feedback control across speech and reaching movements

**DOI:** 10.1101/2021.04.05.438454

**Authors:** Benjamin Parrell, Hyosub E. Kim, Assaf Breska, Arohi Saxena, Richard B. Ivry

## Abstract

Errors that result from a mismatch between predicted movement outcomes and sensory afference are used to correct ongoing movements through feedback control and to adapt feedforward control of future movements. The cerebellum has been identified as a critical part of the neural circuit underlying implicit adaptation across a wide variety of movements (reaching, gait, eye movements, and speech). The contribution of this structure to feedback control is less well understood: although it has recently been shown in the speech domain that individuals with cerebellar degeneration produce even larger online corrections for sensory perturbations than control participants, similar behavior has not been observed in other motor domains. Currently, comparisons across domains are limited by different population samples and potential ceiling effects in existing tasks. To assess the relationship between changes in feedforward and feedback control associated with cerebellar degeneration across motor domains, we evaluated adaptive (feedforward) and compensatory (feedback) responses to sensory perturbations in reaching and speech production in human participants of both sexes with cerebellar degeneration and neurobiologically healthy controls. As expected, the cerebellar group demonstrated impaired adaptation in both reaching and speech. In contrast, the groups did not differ in their compensatory response in either domain. Moreover, compensatory and adaptive responses in the cerebellar group were not correlated within or across motor domains. These results point to a general impairment in feedforward control with spared feedback control in cerebellar degeneration. However, the magnitude of feedforward impairments and potential changes in feedback-based control manifest in a domain-specific manner across individuals.

**Significance Statement:** The cerebellum contributes to feedforward updating of movement in response to sensory errors, but its role in feedback control is less understood. Here, we tested individuals with cerebellar degeneration (CD), using sensory perturbations to assess adaptation of feedforward control and feedback gains during reaching and speech production tasks. The results confirmed that CD leads to reduced adaption in both domains. However, feedback gains were unaffected by CD in either domain. Interestingly, measures of feedforward and feedback control were not correlated across individuals within or across motor domains. Together, these results indicate a general impairment in feedforward control with spared feedback control in CD. However, the magnitude of feedforward impairments manifests in a domain-specific manner across individuals.

## Introduction

Coordinated movement relies on a combination of feedback control and anticipatory mechanisms. A mismatch between the predicted and actual feedback resulting from a motor command can lead to online corrections as well adaptation of feedforward control for future movements (Shadmehr and Krakauer, 2008). The cerebellum plays a critical role in this latter process, helping ensure that the predictive system is optimally calibrated. One line of supportive evidence comes from the substantial literature showing markedly impaired performance of individuals with cerebellar degeneration (CD) during sensorimotor adaptation tasks involving upper limb movement (Martin et al., 1996; Smith and Shadmehr, 2005; Tseng et al., 2007; Donchin et al., 2012; Schlerf et al., 2013), gait (Morton and Bastian, 2006), eye movements (Xu-Wilson et al., 2009), and speech (Parrell et al., 2017).

If, and how, the cerebellum contributes to feedback control is less clear. One clue comes from intentional tremor, a prominent feature of CD where low-frequency oscillations occur around the movement endpoint. Intentional tremor is reduced when movement is produced without visual feedback (Day et al., 1998). Such behavior is broadly consistent with a control system that relies on visual feedback (Beppu et al., 1987), and could occur if the gains on sensory feedback errors are larger in individuals with CD. Larger gains could lead to overcorrections for errors and the need for additional counter-corrections. This hypothesis is supported by evidence showing that, relative to controls, individuals with CD produce larger compensatory responses to auditory perturbations of speech (Parrell et al., 2017; Houde et al., 2019; Li et al., 2019).

There is mixed evidence concerning feedback gains in other types of movement. Individuals with CD produce a smaller long-latency muscle response to mechanical perturbations (Kurtzer et al., 2013), consistent with a decreased gain in the response to proprioceptive feedback. However, their feedback-based corrections during split-belt treadmill walking are relatively normal (Morton and Bastian, 2006), suggesting proprioceptive gains are unaffected for at least some tasks. Moreover, individuals with CD have normal feedback gains in a continuous visual tracking task, but with a substantial phase lag (Zimmet et al., 2020). Computationally, this is consistent with an increased reliance on (delayed) feedback in the absence of a predictive function for state estimation (Wolpert et al., 1998; Miall et al., 2007).

It is possible that potential increases in feedback gains in non-speech tasks may be obscured by a ceiling effect. While the auditory feedback response in speech only partially compensates for the perturbation, the responses to perturbations in tasks such as visually-guided movements or walking typically provide nearly complete compensation to the perturbation (Morton and Bastian, 2006; Tseng et al., 2007). As such, reductions in feedback gains could be readily measured but increases in gain might be hard to detect. Alternatively (or additionally), modification of feedback gains in CD may be task-dependent, with variable changes across movement types. This would stand in contrast to feedforward control, where the cerebellum appears to play a similar role in implicit adaptation across domains.

The preceding hypotheses are based on inferences drawn from disparate studies that distinct methods and typically focus on a single motor domain. In the present study, we used a 2 × 2 design to evaluate feedback and feedforward control in two motor tasks, one involving visually-guided reaching and the other speech production. To avoid potential ceiling effects in the former, we used a task shown previously to induce only partial corrections to the visual perturbation (Körding and Wolpert, 2004). Moreover, by testing the same individuals on all four tasks, we can perform a correlational analysis, focusing on two key questions related to individual differences associated with CD. First, are patterns of impairment similar across the two motor domains and/or forms of control? Second, is the degree of impairment in feedforward control (adaptation) predictive of feedback gains, a signature that would be consistent with the hypothesis that enhanced feedback gains arise as a compensatory mechanism.

## Materials and Methods

### Participants

23 individuals with cerebellar degeneration (CD: 19 female, 37-89 years, mean age 62 years) and 15 age-matched neurobiologically healthy controls (CO: 8 female, 43-85 years, mean age 61 years) were recruited for the study. One CD participant was excluded due to a potential mild cognitive impairment (see below), leaving 22 individuals in the CD group. The participants had normal or corrected-to-normal vision, and none reported any history of speech or hearing impairments, or significant neurological issues (other than ataxia in the CD group). The participants were naïve as to the experimental hypotheses and none had participated in our previous experiment using altered auditory feedback (Parrell et al., 2017). Participants provided informed consent and received financial compensation for their time. The protocol was approved by the institutional review boards at the University of Wisconsin–Madison and the University of California, Berkeley.

The patients were drawn from the Northern California Ataxia Support group, the Madison Ataxia Support group, the Kansas City Ataxia Support group, and individuals who attended the 2018 annual meeting of the National Ataxia Foundation. Recruitment for the latter two groups was conducted in coordination with officials of the associated organizations. In terms of etiology, the CD group was heterogenous: 11 had a confirmed genetic subtype (3 SCA3; 6 SCA6; 2 SCA8), 1 had a clinically-diagnosed genetic subtype (AOA 2), and 11 had been diagnosed as cerebellar ataxia of unknown etiology. Inclusion was limited to individuals who had symptoms of ataxia in combination with genetic confirmation, radiological findings of cerebellar atrophy, or medical diagnosis indicating cerebellar atrophy. While we performed a brief neurological exam to assess ataxia (see below), the other information was based on the participants’ self-report. We note that individuals who attend support groups are generally very proactive and well-informed about their condition. Age-matched control participants were drawn from the greater Berkeley community.

All of the participants were administered the Montreal Cognitive Assessment (MoCA) as a gross measure of cognitive impairment. The CD group was additionally administered the Standard Ataxia Rating Scale (SARA) to assess the severity of their ataxia. SARA subscores were calculated for both upper limb control and speech. We did not include a formal assessment of signs of extracerebellar degeneration, although none of the participants, including those with SCA3, exhibited gross symptoms of basal ganglia dysfunction such as rigidity or resting tremor. One individual with CD was excluded due a MOCA score < 18, indicative of moderate cognitive impairment. See the Table 1 for a full characterization of the CD group.

**Table 1:** Characteristics of individuals with cerebellar degeneration. One participant, shown in gray, was excluded due to a MOCA score indicative of moderate cognitive impairment.

| Diagnosis | Age | MOCA | SARA overall score | SARA upper limb subscore | SARA intention tremor subscore | SARA speech subscore |
| --- | --- | --- | --- | --- | --- | --- |
| SCA3 | 61 | 28 | 16 | 5 | 1.5 | 3 |
| SCA3 | 75 | 26 | 16.5 | 5.5 | 1.5 | 0 |
| SCA3 | 65 | 25 | 24 | 7 | 2 | 1 |
| SCA6 | 77 | 22 | 8 | 2 | 1 | 1 |
| SCA6 | 70 | 23 | 7.75 | 3.25 | 0.5 | 0 |
| SCA6 | 60 | 26 | 3 | 0.5 | 0 | 0 |
| SCA6 | 49 | 26 | 1 | 1 | 0 | 0 |
| SCA6 | 65 | 24 | 9 | 3.5 | 0.5 | 0 |
| SCA6 | 57 | 27 | 6 | 3 | 0 | 0 |
| SCA8 | 57 | 27 | 11 | 3.5 | 1 | 0 |
| SCA8 | 53 | 30 | 3.5 | 2 | 0 | 0 |
| AOA2 | 42 | 27 | 22 | 6 | 2.5 | 2 |
| CD, unknown | 57 | 22 | 10 | 4 | 1 | 2 |
| CD, unknown | 65 | 17 | 25 | 6 | 2 | 3 |
| CD, unknown | 89 | 26 | 9.5 | 2 | 0.5 | 4 |
| CD, unknown | 68 | 27 | 23 | 5 | 2 | 1 |
| CD, unknown | 64 | 25 | 8 | 1.5 | 0.5 | 1 |
| CD, unknown | 37 | 27 | 14 | 3 | 1 | 2 |
| CD, unknown | 72 | 27 | 18.5 | 4 | 1.5 | 1 |
| CD, unknown | 64 | 28 | 11 | 3.5 | 0 | 1.5 |
| CD, unknown | 54 | 28 | 8.5 | 4.5 | 0.5 | 3 |
| CD, unknown | 65 | 25 | 9.5 | 2.5 | 1 | 0 |
| CD, unknown | 62 | 30 | 12 | 5.5 | 1 | 1 |

Data from one control participant was excluded from Exp 4 due to equipment failure during data collection. The final sample size was based on typical sample sizes in previous studies (e.g., Morton and Bastian, 2006; Kurtzer et al., 2013; Parrell et al., 2017; Zimmet et al., 2020). Given the final sample sizes of 22 and 15 for the CD and control groups, respectively, we have a power of 0.75 to detect a large between group effect (*d* = 0.8). The larger sample for the CD group has a power of 0.75 to detect a medium effect (*d* = 0.5) for the within-group correlations. Note that we tested a larger group of individuals in the CD group to have sufficient power to examine predicted correlations within this group of the behavioral measures and clinical symptoms and, more important, a critical prediction concerning the relationship of feedforward and feedback control.

### Experimental Design

Each participant completed four conditions in a single experimental session that lasted approximately one hour, including breaks of approximately 5 min between each condition. Feedforward and feedback control during reaching were assessed in Conditions 1 and 2, respectively; similarly, feedforward and feedback control during speech were assessed in Conditions 3 and 4, respectively. A schematic of each of the four conditions is shown in Figure 1. The same order of the four conditions was used for each participant. While this approach introduces order confounds, we opted to keep the order fixed given that a) this is preferable for correlational analyses across tasks, as counterbalancing task order introduces additional between-participant variance to these analyses (Goodhew and Edwards, 2019) and b) the sample size was insufficient to assess effects of task order in a counterbalanced design. Stimulus presentation and data collection was controlled with Matlab for all conditions.

**Figure 1:**
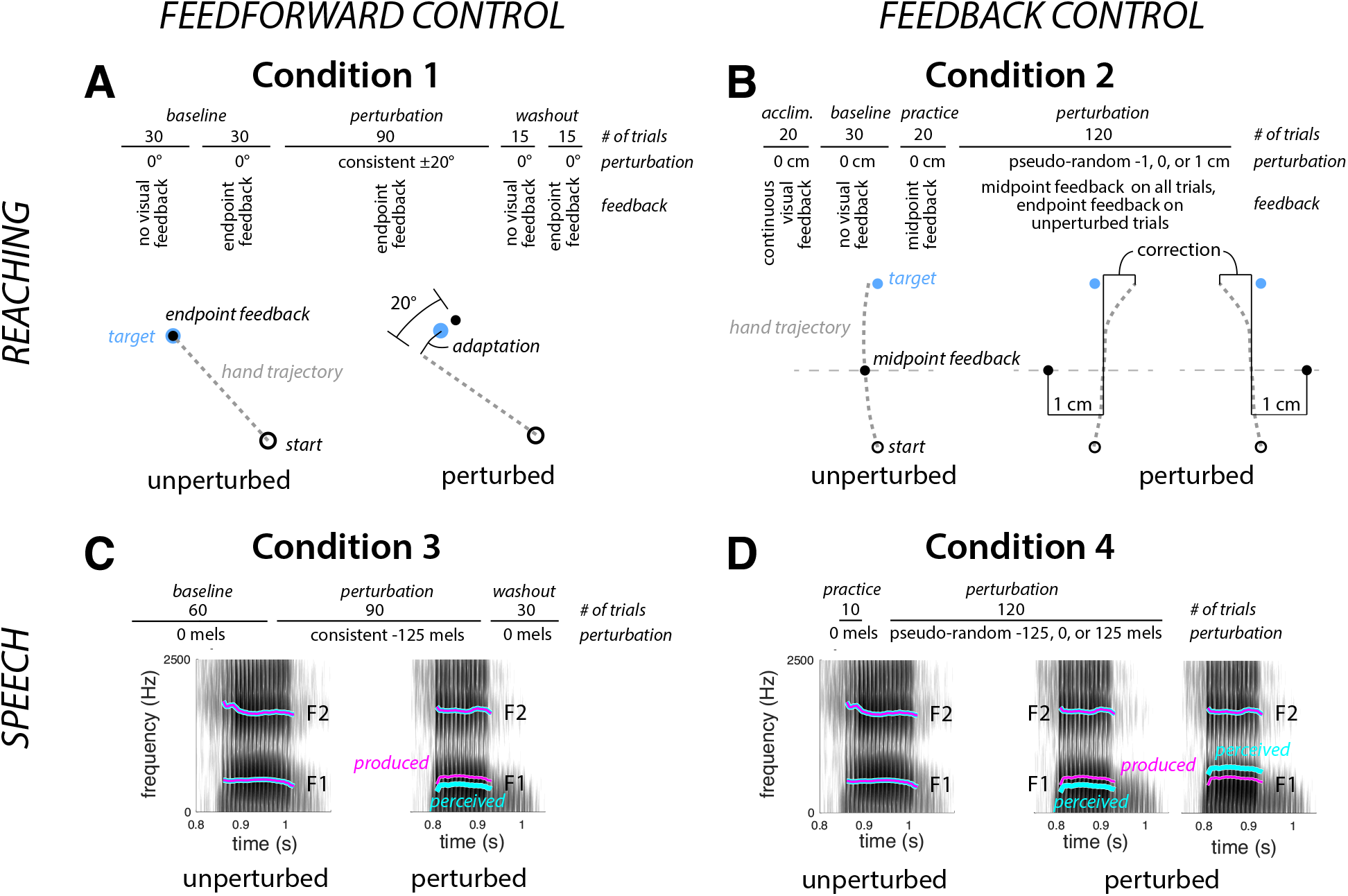
Experimental design. The top row of each panel shows the structure of the experimental session and the bottom part depicts the perturbation. **A:** Condition 1, feedforward control in reaching: Participants made 8 cm center-out reaches to targets (blue dot) with endpoint feedback. During the perturbation phase, the location of the feedback cursor was rotated by 20° from actual hand position (direction cross-balanced across participants). The dashed line represents the unseen hand movement. **B:** Condition 2, feedback control in reaching: Participants made 16 cm reaches to a target (blue dot). During the perturbation phase, visual feedback about hand position (black dot) was given at reach midpoint with the feedback shifted by -1, 0, or 1 cm, with the shift on each trial determined in a pseudo-random manner. The thin gray line depicts location of reach midpoint, but was not visible to the participant. **C:** Condition 3, feedforward control in speech: Participants spoke a single word on each trial, hearing a playback of their speech over headphones with minimal delay. During the perturbation phase, feedback of the first formant during the vocalic portion of the utterance was perturbed by imposition of a – 125 mel shift (cyan trace superimposed on the spectrogram). **D:** Condition 4, feedback control in speech: Participants spoke a single word on each trial. During the perturbation phase, auditory perturbations (cyan trace) of -125, 0, or 125 mels were pseudo-randomly applied to the first formant during the vocalic portion of the utterance.

#### Condition 1: Feedforward control in reaching

Participants were seated in front of a 53.2 × 20 cm LCD screen (ASUS) that was horizontally encased in a table frame mounted 27 cm above a 49.3 × 32.7 cm digitizing tablet (Intuos 4XL, Wacom, Vancouver, WA). Participants held a modified air hockey paddle and made reaching movements by sliding the paddle across the table. The position of a stylus embedded in the paddle was recorded by the tablet at 200 Hz. Feedback, when available, was presented in the form of a cursor on the LCD screen. Participants’ view of their hand was blocked by the LCD screen. To further limit vision of the upper arm, the experiment was conducted in a darkened room. The experiment was controlled with Psychtoolbox (Brainard, 1997; Pelli, 1997; Kleiner et al., 2007).

Participants made center-out reaches, moving to targets located at a radial distance of 8 cm from the center of the workspace. The start location was marked by a white ring (6 mm diameter). On each trial, the participant moved the digitizing stylus to position the hand within the start location. Visual feedback about the stylus position was provided by a small (3.5 mm diameter) white circle. After the participant had maintained the stylus position within the start location for 500 ms, one of 3 equally-spaced targets (6 mm diameter) appeared (0°, 120°, or 240°). The participant was instructed to reach, attempting to “slice through” the target. The instructions emphasized that the movements should be made quickly and, to minimize demands on endpoint (radial) accuracy, should terminate beyond the target location. RT was not emphasized; the movement could be initiated at any time after the presentation of the target.

Visual feedback about the movement was limited to endpoint feedback, eliminating the opportunity for visually guided corrections. The feedback cursor disappeared at movement onset and reappeared when the radial distance of the hand movement reached 8 cm. The cursor remained visible for 50 ms and then was blanked. If participants reached the 8 cm threshold in ≤ 500 ms, a knocking sound was played, indicating that the movement was “fast enough and far enough.” If the movement duration was > 500 ms, participants heard a recording of the words “too slow”. During the return movement to the start location, visual feedback was withheld until the stylus was within 3 cm of the start position.

The condition consisted of five phases. 1) A *no feedback baseline* phase of 30 trials to measure movement variability in the absence of visual feedback. 2) A *baseline phase* of 30 trials with veridical endpoint feedback of the cursor. 3) A *perturbation phase* of 90 trials in which the location of the cursor at movement endpoint was rotated 20° from the true position. The 20° rotation was chosen to limit awareness of the perturbation and minimize the use of explicit reaiming (Bond and Taylor, 2015; Morehead et al., 2015; Werner et al., 2015). The perturbation was either clockwise or counter-clockwise, counterbalanced across participants. 4) An *aftereffect phase* of 15 trials in which there was no visual feedback. 5) A *washout* phase of 15 trials with veridical endpoint feedback.

The hand angle for each trial was measured as the angle between a line connecting the start location and the hand position at the time of peak radial velocity, relative to the line connecting the start and target locations. To remove intrinsic biases in reaching, the mean hand angle during the baseline phase was subtracted from the heading angle for each trial. We focus on two measures of adaptation, the mean of the last 15 trials of the perturbation phase (*asymptote*) and the mean of the 15 trials in the aftereffect phase. We additionally measured reaction time (time from target appearance to movement onset) and movement time (time from movement onset to target), as well as the percentage of trials where any part of the cursor representing hand position overlapped with the target (*hits*).

#### Condition 2: Feedback control in reaching

The experimental apparatus was identical to Exp 1. On each trial, the white ring indicating the start location appeared near the bottom edge of the screen at the horizontal meridian. After this positioned was maintained for 500 ms, the target appeared at a fixed position, 16 cm in front of the start location. The longer distance was used so that feedback could be presented at the midpoint of the movement, providing sufficient distance (and time) for an online correction.

There were five phases to Condition 2. 1) An *acclimation phase* of 20 trials with continuous veridical feedback. 2) A *baseline phase* of 30 trials in which there was no visual feedback, either during the movement or at the target distance. 3) A *practice* phase of 20 trials to introduce the method for providing limited visual feedback. The cursor appeared twice on these trials, first when the radial amplitude of the stylus was 8 cm from the start location (midpoint, duration = 100 ms), and second when the radial amplitude reached 16 cm (endpoint, duration = 50 ms). Participants were instructed to “use what you see midway through the reach to get as close as possible to the target”. 4) A *perturbation phase* of 120 trials with midpoint feedback. On 50% of the trials, the midpoint feedback was given at the true location of the stylus, and on the remaining 50% of the trials, the midpoint feedback was shifted 1 cm to the left of the stylus position or 1 cm to the right (25% each). Each cluster of 4 reaches consisted of 2 unperturbed trials, and 1 trial with the leftward and rightward perturbation, in a randomized order. Endpoint feedback was not provided on perturbed trials to minimize learning effects (i.e., anticipatory effects based on prior responses to the perturbation). Endpoint feedback was provided on the unperturbed trials to help participants remain calibrated to the target reaching location. 5) A *final phase* of 20 trials with no visual feedback. Data from this last phase was not analyzed further.

As in Condition 1, participants were instructed to slice through the target. The movement time criterion was increased to 1200 ms. The extra time was employed to compensate for the larger amplitude movements and to encourage participants to adopt a movement speed that allowed them time to use the midpoint feedback to make an online correction (if warranted). The knocking sound was played if the target amplitude was reached within 1200 ms, and if the movement duration was between 400 and 1200 ms, the target circle turned green. If the movement was < 400 ms, the words “too fast” appeared on the screen. If the movement duration was longer than 1200 ms, the target circle turned red and a recording of the words “too slow” was played.

The focus here was on the online corrections made in response to the horizontally displaced midpoint feedback; that is, the horizontal position of the hand relative to the target at the target distance. However, the raw horizontal displacement would also reflect the horizontal position of the hand at reach midpoint, which may vary between trials. To account for this, we fit the trial-by-trial data of each participant with a linear model that predicted the final horizontal position of the hand on each trial from the horizontal hand position at reach midpoint, the visual perturbation (treated as a categorical variable and coded using separate dummy predictors for each direction), and the interaction between midpoint hand position and the perturbation:

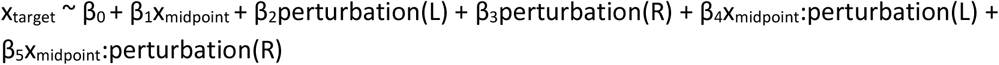

This model allows us to estimate two dependent measures of feedback-based corrections: 1) the magnitude of the correction for the perturbation (direction effect estimates, β_2_, β_3_) and 2) corrections for self-produced variability (the midpoint position estimate, β_1_, and its adjustment in the presence of perturbations, β_4_, β_5_). For statistical analysis, the sign of β_2_ values were flipped such that negative values always reflected a compensatory response to the perturbation, regardless of perturbation direction. Additionally, this method allows us to estimate the effect of the correction for the visual perturbation while accounting for any individual differences in the bias of the overall reach trajectory, an effect captured by the intercept term (β_0_), which reflects the horizontal displacement relative to the target position for unperturbed reaches. We measured movement time, reaction time, and proportion of trials where the cursor hit the target as for Condition 1. Hit proportion was calculated only for unperturbed trials.

#### Condition 3: Feedforward control in speech

Participants were again seated in front of the horizontally-aligned monitor. Participants wore a head-mounted microphone (AKG C520) and close-backed, over-the-ear headphones (Beyerdynamic DT 770). On each trial, participants spoke the word “head” after it appeared on the monitor. The utterance was digitized through a Scarlett 2i2 sound card and processed with Audapter (Cai et al., 2008; Tourville et al., 2013) to synthesize a sound for playback over the headphones. The recording, processing, and playback occurred in near real time (∼20 ms delay). The intensity of the synthesized feedback was set to roughly 80 dB based on each participants comfortable speaking volume, mixed with speech-shaped noise at ∼60 dB to mask air- or bone-conducted direct feedback of the utterance. The actual intensity level of the feedback varied with changes in the intensity of participants’ speech.

There were four phases in Condition 3. For each phase, the word “head” remained visible for 2.5 s on each trial, and the trials were separated by 1.1-1.3 s (randomly jittered). 1) A *baseline phase* of 60 trials with speech feedback resynthesized through Audapter with no auditory perturbation applied. 2) A *perturbation phase* of 100 trials during which a constant shift of -125 mels was applied to the first vowel formant (F1) throughout the utterance, shifting the F1 value of “head” towards that of “hid” (<u>mels are a perceptually-calibrated unit of frequency where each step is perceived as equivalently distinct across all frequencies, Stevens et al., 1937</u>). 3) A *washout phase* of 30 trials in which there was no auditory perturbation.

Vowel onset and offset was labelled for each trial using a semi-automated procedure. First, automatic labels were generated by identifying on the waveforms where the speech amplitude first crossed (onset) or fell below (offset) a participant-specific amplitude value. The automatic labels were then visually inspected and corrected when the waveform and spectrogram indicated that the automatic markings were inappropriate. A semi-automated procedure was used to track the formants during the vocalic phase of the utterance using participant-specific vales for LPC order and pre-emphasis. Formant tracking was performed with Praat (Boersma and Weenink, 2019). Using the Wave Runner software package (Niziolek and Houde, 2015), the tracks were visually inspected and, when they did not align with visible formants on the speech spectrogram, the formant tracking parameters (pre-emphasis, LPC order) were modified.

The primary outcome measure was adaptation of the vocalic portion of the utterances across trials. For each trial, we calculated the average F1 value from 50-100 ms after vowel onset. This window avoids the transitional phase of the formant during the word-initial consonant (initial 50 ms) as well as any online compensatory response to the perturbation, which typically begins later than 100 ms after vowel onset (Tourville et al., 2008; Cai et al., 2012; Parrell et al., 2017). To facilitate comparisons across participants, the F1 values were normalized with respect to the average F1 value taken over the 50-100 ms window during the second half of the baseline phase (trials 31-60). We chose to use the second half of the baseline phase to allow participants time to acclimate to the processed auditory feedback.

#### Condition 4: Feedback control in speech

The stimulus set consisted of four words, “dead”, “fed”, “said”, and “shed”, selected because they share the same vowel /ε/. The condition began with a calibration phase designed to shape the participant’s speaking rate such that the produced vowel duration would be between 300 and 500 ms. This criterion was important to ensure that there would be sufficient time for feedback-based corrections, shown in previous work to have a latency of ∼150 ms (Tourville et al., 2008; Cai et al., 2012; Parrell et al., 2017). On each trial, one of the four words was displayed. After each utterance, the automated estimate of the vowel duration in milliseconds was displayed on the monitor. This procedure was repeated for 10 trials. If the duration fell outside the 300-500 ms window on more than 2 of these 10 trials, the procedure was repeated for 10 more trials. When the utterances fell within the criterion window for at least 8 of the 10 trials, the main experiment began.

Condition 4 had two phases. 1) A *practice phase* of 10 trials with veridical auditory feedback. 2) A *perturbation phase* of 120 trials. During the perturbation phase, participants heard veridical feedback on 50% of trials, a + 125 mel shift of F1 (moving the vowel towards that in “had”) on 25% of trials, and a -125 mel shift of F1 (moving the vowel towards that in “hid”) on the remaining 25% of trials. Each group of 4 trials consisted of 2 unperturbed trials, and 1 trial each of positive and negative F1 perturbations, randomly ordered. Stimuli were presented in a random order (selection with replacement), with the constraint that the total number of each stimulus word be as equal as possible across the experimental condition.

Vowels were tracked as in Exp 3. The primary outcome measure was the online correction to the perturbation, calculated following standard approaches (Cai et al., 2012; Niziolek and Guenther, 2013; Parrell et al., 2017; Daliri et al., 2020). First, an average baseline F1 trajectory was calculated for each stimulus word from the unperturbed productions of that word. Second, the F1 trajectory from each perturbed trial was normalized by subtracting the appropriate word-specific average baseline F1 trajectory, giving a normalized F1 response for each perturbed trial. Trajectories from trials with upward and downward F1 perturbations were separately averaged to generate an average F1 response in each direction. To generate a composite feedback response, the sign of the average response to the upward perturbation was flipped, such that positive values always reflected a compensatory response to the perturbation, regardless of perturbation direction. Note that because of the normalization process (and lack of an explicit target), this differs slightly from the approach in Exp 2, where midpoint variability had to be accounted for. To quantify the magnitude of the compensatory response, the mean value of each average F1 response trajectory between 200-300 ms after vowel onset was calculated. This window begins well after the expected 150 ms latency of the compensatory response is ∼150 ms (Tourville et al., 2008; Cai et al., 2012; Parrell et al., 2017).

### Statistical Analysis

Statistical analysis for all conditions was conducted in R (R Core Team, 2020) using mixed ANOVAs or Welches’ two-sample t-tests. When necessary, post-hoc tests were conducted with a Tukey HSD correction. Where appropriate, one-sample t-tests with Holm-Bonferroni corrections were used to determine whether the responses for each group differed significantly from 0. Means are reported with standard deviations. Effect sizes are given as Hedges *g* or partial eta-squared.

For conditions 1 and 3, a mixed ANOVA was used that included group (ataxic vs control) and phase (end of perturbation and aftereffects), as well as the interaction between the two factors. Similarly, for condition 4, the mixed ANOVA included group (ataxic vs control) and perturbation direction (up and down), as well as the interaction between the two.

For condition 2, separate ANOVAs were used to evaluate differences between the ataxic and control group in 1) the magnitude of the correction in response to the perturbation and 2) the magnitude of correction in response to self-produced variability. These were conducted as second-level analyses, on the beta coefficients estimated in the first-level analyses that were conducted within each individual (see above). The dependent variable for the perturbation correction model was the direction effect coefficient, estimated separately for each perturbation direction (β_2_, β_3_). In addition to the perturbation direction factor (left vs right), this model included a group factor as well as the interaction between perturbation direction and group. For the variability correction, the dependent variable was the midpoint coefficient representing the magnitude of correction for variability at reach midpoint in unperturbed (β_1_), left (β_1_+β_4_), and right (β_1_+β_5_) conditions. In addition to the main variable of interest, group, this model also included perturbation condition (no perturbation, left, right). Welches’ t-tests were used to compare the ataxic and control groups on the reaction time, movement time, and hit percentage in unperturbed trials.

In order to assess the relationship between feedback and feedforward control, we conducted a set of planned correlational analyses using data from pairs of conditions. One set of correlations compared the measures of feedback and feedforward control within each motor domain (i.e., one correlation between the two reach tasks and a second between the two speech tasks). We also performed a second set of correlations of similar forms of control across the motor domains (i.e., one correlation of the feedforward measures from the reach and speech tasks, and a second correlation of the feedback measures from the reach and speech tasks). Because the primary aim of these analyses is to investigate the possibility that changes in motor behavior due to cerebellar degeneration are correlated across sensory domains and control systems, we limited these correlational analyses to the data from the CD group. We also examined the extent to which our experimental measures of feedforward and feedback control in the ataxic group were related to ataxia symptom severity, as measured by SARA speech and upper limb subscores.

## Results

### Condition 1: Feedforward control in reaching

In the baseline phase with visual feedback, the CD group was substantially more variable than controls (standard deviation of 6.2 ± 2.0° vs 4.0 ± 0.8°, t(31.6) = 4.6 p < 0.0001, g = -0.89) and less accurate (percentage of targets hit, 67 ± 25% vs 87 ± 15%, t(35.8) = -3.0, p = 0.005, g = 1.28). The CD group also produced slower movements than controls (347 ± 148 ms vs 264 ± 61 ms, t(31.6) = 2.4, p = 0.02, g = 0.67) and had longer reaction times (544 ± 116 ms vs 442 ± 58 ms, t(34.0) = 3.6, p = 0.001, g = 1.02).

Our principle outcome measure, adaptation, was operationalized as the change in heading angle following the introduction of a 20° rotation of the feedback cursor. As can be seen in Figure 2, the perturbation induced a gradual change in heading angle, with the functions appearing to reach or approach asymptote by the end of the perturbation trials. Adaptation in the control participants compensated for approximately 83% of the perturbation; the comparable figure for the CD participants was only 45%. A decline in adaptation is visible throughout the aftereffect phase in which there was no visual feedback and the following washout phase in which veridical feedback was reintroduced. In Figure 2 and subsequent figures, red diamonds are used to indicate individuals with SCA6 and SCA8, two genetic subtypes in which the degeneration is largely confined to the cerebellum proper; red circles indicate CD individuals with other diagnoses. As can be seen, the relatively “pure” cerebellar participants fall within a similar distribution as the other CD participants.

**Figure 2:**
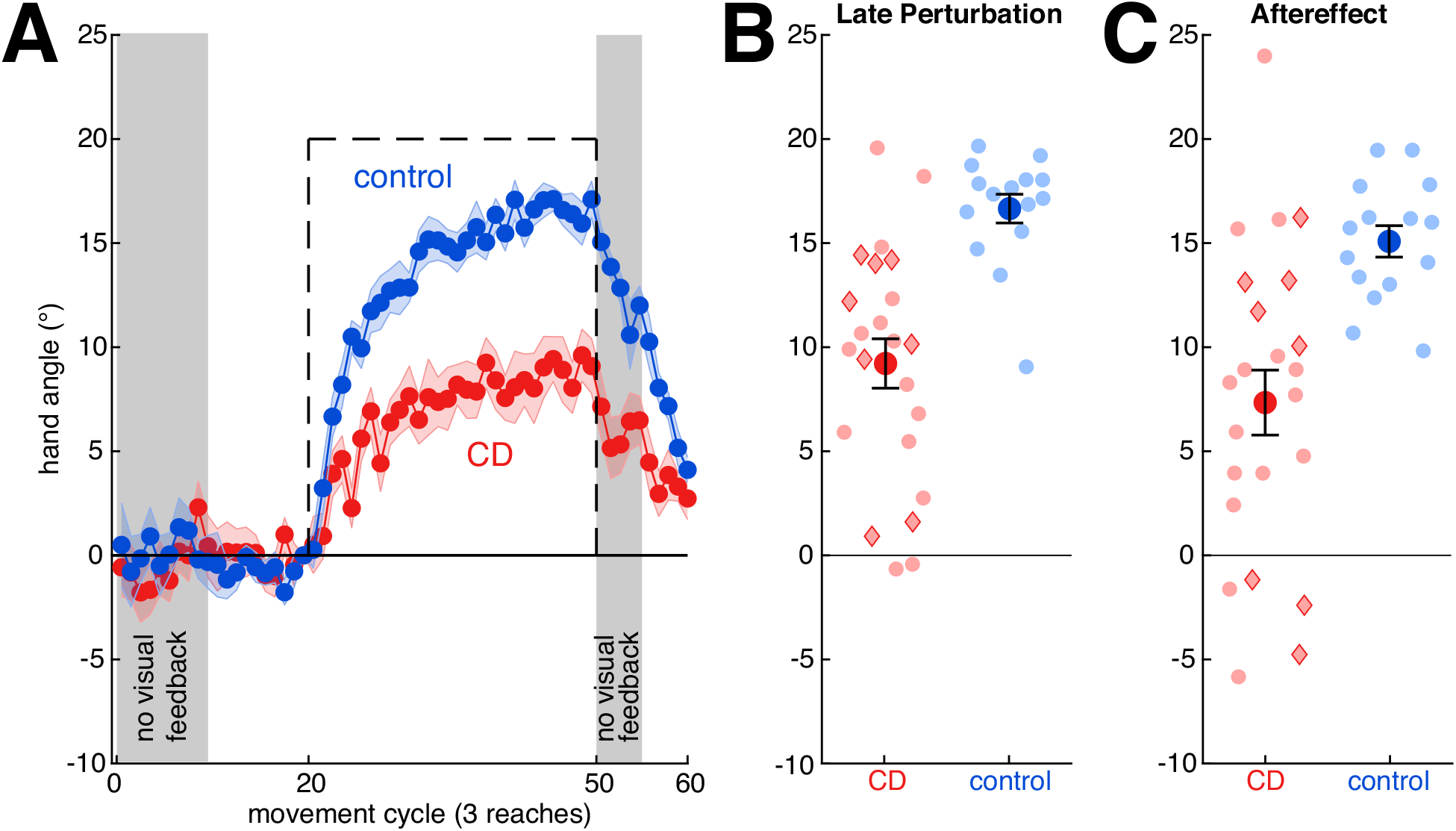
In the visuomotor rotation task, the CD group reached a lower level of adaptation and exhibited a smaller aftereffect relative to the control group. **A:** Hand angle over the course of the experimental condition. Means and standard errors are shown for control participants (blue) and individuals with cerebellar degeneration (red). Visual feedback is withheld in the shaded phases. The perturbation is shown with a dashed black line. **B**,**C:** Individual data (semi-transparent dots) and group means ± standard error for the CD and control participants during the last 15 reaches of the perturbation phase (B) and 15-trial aftereffect phase in which the visual feedback was withheld (C). Red diamonds indicate individuals with SCA6 and SCA8; red circles indicate the other CD participants.

Although we did not conduct any post-session interviews, we assume that adaptation was implicit with negligible contributions from strategic aiming (e.g., in the opposite direction of the perturbation). Not only does the size of the perturbation fall within a range in which learning is thought to be driven largely by implicit adaptation (Bond and Taylor, 2015; Morehead et al., 2015; Werner et al., 2015), but there is only a slight decrease in hand angle at the start of the aftereffect block. Moreover, we did not observe a sustained increase in reaction times following the introduction of the perturbation, a cardinal signature of aiming (McDougle and Taylor, 2019), in our data (comparing first 10 trials of the perturbation to the baseline, *p* > 0.94 for both groups).

Statistically, we first confirmed that the sign-dependent changes in heading angle were significantly different than zero, the signature of adaptation. When measured during the final trials of the perturbation block, the heading angle values were different than zero for both the control (16.6 ± 2.7°, *t*(14) = 24.1, *p* < 0.0001, *g* = 5.90) and CD groups (9.2 ± 5.7°, *t*(22) = 7.8, *p* < 0.0001, *g* = 1.56). The adapted response persisted into the aftereffect block (controls: 15.1 ± 2.9°, *t*(14) = 20.0, *p* < 0.0001, *g* = 4.87; CD: 7.3 ± 7.5°, *t*(22) = 4.7, *p* < 0.0001, *g* = 0.95), although the value was smaller than observed at the end of the perturbation block for both groups (*F*(1,36) = 7.4, *p* = 0.01, *η*^*2*^ = 0.17). Across both the perturbation and aftereffect phases, adaptation was significantly greater in the control group (*F*(1, 36) = 20.1, *p* < 0.0001, *η*^*2*^ = 0.36) and the Group x Phase interaction was not significant (*F*(1,36) = 0.05, *p* = 0.82, *η*^*2*^ = 0.001). These results are in accord with previous results (for a review, see Krakauer et al., 2019) in showing that adaptation of feedforward control for reaching is impaired in individuals with CD.

### Condition 2: Feedback control during reaching

As in Condition 1, there were some kinematic differences between the two groups. The most salient group difference was in movement accuracy: control participants hit the target more often than the CD participants on the unperturbed trials (control: 85 ± 10 %, CD: 70± 24%, t(31.9) = 2.7, *p* = 0.01, *g* = 0.75). There were small differences between the groups in reach curvature, with controls showing a slight leftward shift from midpoint to target on unperturbed trials and the CD group showing a slight rightward shift on these trials (control: -0.17 ± 0.193 cm, CD: 0.13 ± 0.52 cm, *t*(30.2) = 2.5, *p* = 0.02, *g* = 0.69, **Figure 3**F). Reaction times were also slower in the CD group (control: 452 ± 79 ms, CD: 525 ± 138 ms, *t*(35.5) = 2.08, *p* = 0.045, *g* = 0.61) whereas movement time was similar between groups (control: 717 ± 114 ms, CD: 722 ± 114 ms, *t*(30.1) = 0.14, *p* = 0.89, *g* =0.05).

**Figure 3:**
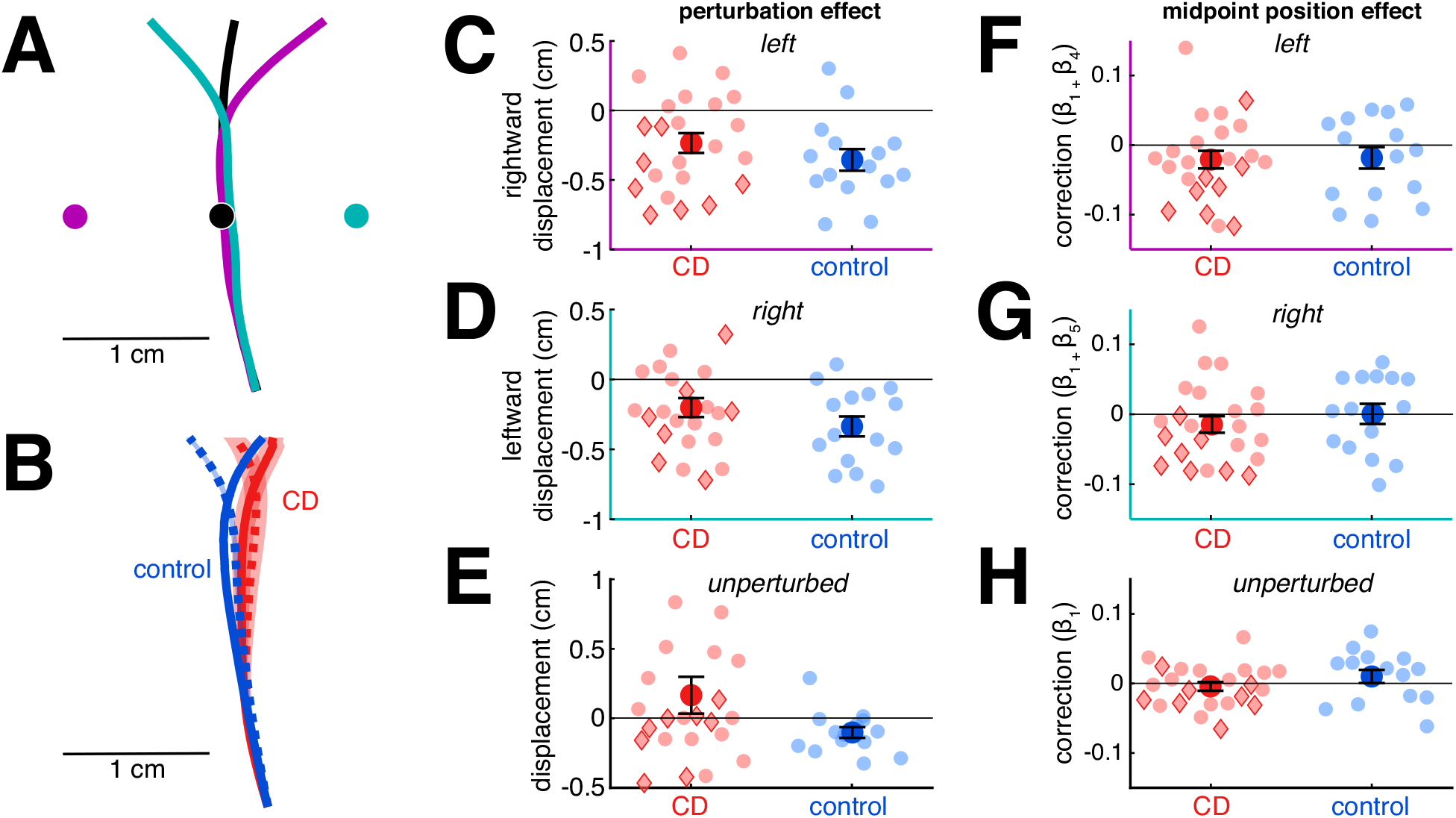
In the reaching feedback control task, both the CD and control groups showed a robust online correction to perturbed visual feedback presented at the midpoint of the reach, with no group differences. **A:** Example data from one control participant showing average reach trajectories for unperturbed reaches (black), reaches with a 1 cm rightward visual perturbation (teal), and reaches with a 1 cm leftward visual perturbation (purple). Dots in corresponding colors show the location of the visual feedback. The corrective response only partially corrects for the perturbation. **B**: Reach trajectories for the CD group (red) and control group (blue) aligned to the same starting point, showing mean responses to leftward perturbations (solid lines) and rightward perturbations (dashed lines). Shaded areas represent standard error across participants. **C, D**: Compensatory response to the feedback perturbation. **E:** Estimate of the change in horizontal hand position from reach midpoint on unperturbed trials. **F, G, H:** Estimates of the effect of true hand position at reach midpoint on the compensatory response for leftward (**F**), rightward (**G**), and unperturbed (**H**) trials. Across all plots, compensatory responses have negative values. Semi-transparent dots represent individuals. Red diamonds indicate individuals with SCA6 and SCA8; red circles indicate the other CD participants.

Our principle dependent measure, online corrective responses, was operationalized as a lateral shift in the trajectory in response to perturbed feedback that was presented at the midpoint of the movement, with the direction of the perturbation randomized across trials. As can be seen in the data from a representative control participant (Fig 3A), the hand trajectory deviated in the opposite direction of the perturbation, a signature of a feedback-based response. The lateral shift accounted for roughly 33% of the perturbation magnitude in the control group and 21% in the ataxic group (Fig 3B, D). These corrective feedback responses were significantly different from 0 in both groups in response to the leftward perturbation (CD: -0.21 ± 0.34 cm, *t*(22) = 3.3, *p* = 0.003, *g* = 0.67; control: -0.35 ± 0.30 cm, *t*(14) = 4.6, *p* = 0.0005, *g* = 1.12) and rightward perturbation (CD: -0.20 ± 0.32 cm, *t*(22) = 2.9, *p* = 0.007 *g* = 0.60; control: -0.34 ± 0.28 cm, *t*(14) = 4.7, *p* = 0.0003, *g* = 1.15). While the magnitude of the feedback-based response was smaller for the CD group, this difference was not significant (*F*(1,36) = 2.8, *p* = 0.1, *η*^*2*^ = 0.07). There was no difference between the responses to the two perturbation directions (*F*(1,36) = 0.16, *p* = 0.69, *η*^*2*^ = 0.004) and the Group x Direction interaction was not significant (*F*(1,36) = 0.009, *p* = 0.93, *η*^*2*^ = 0.0002). These results indicate that feedback-based corrective responses to visual perturbations are not enhanced in individuals with CD; indeed, as a group the trend was for attenuated feedback responses.

In a second measure of feedback control, we looked at adjustments in the reach trajectory in response to motor variability (estimated as the change in horizontal position from reach midpoint to endpoint unrelated to the visual perturbation; model parameters β_1_, β_1_+β_4_, and β_1_+β_5_). Although both groups showed a slight shift in trajectory in the compensatory direction in response to the position of the cursor at reach midpoint (**Error! Reference source not found**.C,E,G), this shift was not significantly different from 0 after correction for multiple comparisons (CD: -0.010 ± 0.035, p = 0.05, *g* = -0.30 ; control: -0.009 ± 0.042, p = 0.24, *g* = - 0.21). There was no difference between the two groups (*F*(1,36) = 0.01, *p* = 0.91, *η*^*2*^ = 0.0004), nor any interaction between group and direction (*F*(2,72) = 2.4, *p* = 0.10, *η*^*2*^ = 0.06). However, there was a significant effect of perturbation direction (*F*(2,72) = 7.3, *p* = 0.001, *η*^*2*^ = 0.17) such that the shift in trajectory was greater in the presence of leftward perturbations (−0.021 ± 0.0332) compared to trials with no perturbation (0.001 ± 0.038; *p* = 0.008, *g* = 0.62). There was a similar trend in for trials with rightward perturbations (−0.010 ± 0.036), but this did not reach significance (*p* = 0.15, *g* = 0.33). In sum, there is little evidence that either group corrected for self-produced variability at reach midpoint.

### Condition 3: Feedforward control in speech

Adaptation was operationalized as the change in F1 in response to a -125 mel F1 shift in the feedback heard by participants, introduced via the real-time resynthesis of their utterances. As can be seen in Figure 4A, this auditory perturbation caused a gradual change in F1 that opposed the perturbation. At asymptote, the adaptive response had reached approximately 40% of the perturbation in the control group and 25% in the CD group. The lower degree of compensation relative to reaching is consistent with previous studies (Houde and Jordan, 1998; Purcell and Munhall, 2006; Parrell et al., 2017). Note that, unlike Condition 1, post-perturbation trials (washout phase) always included veridical feedback; we did not include a no-feedback phase as masking auditory feedback with noise leads to substantial changes in speech (Lombard, 1911; Summers et al., 1988).

**Figure 4:**
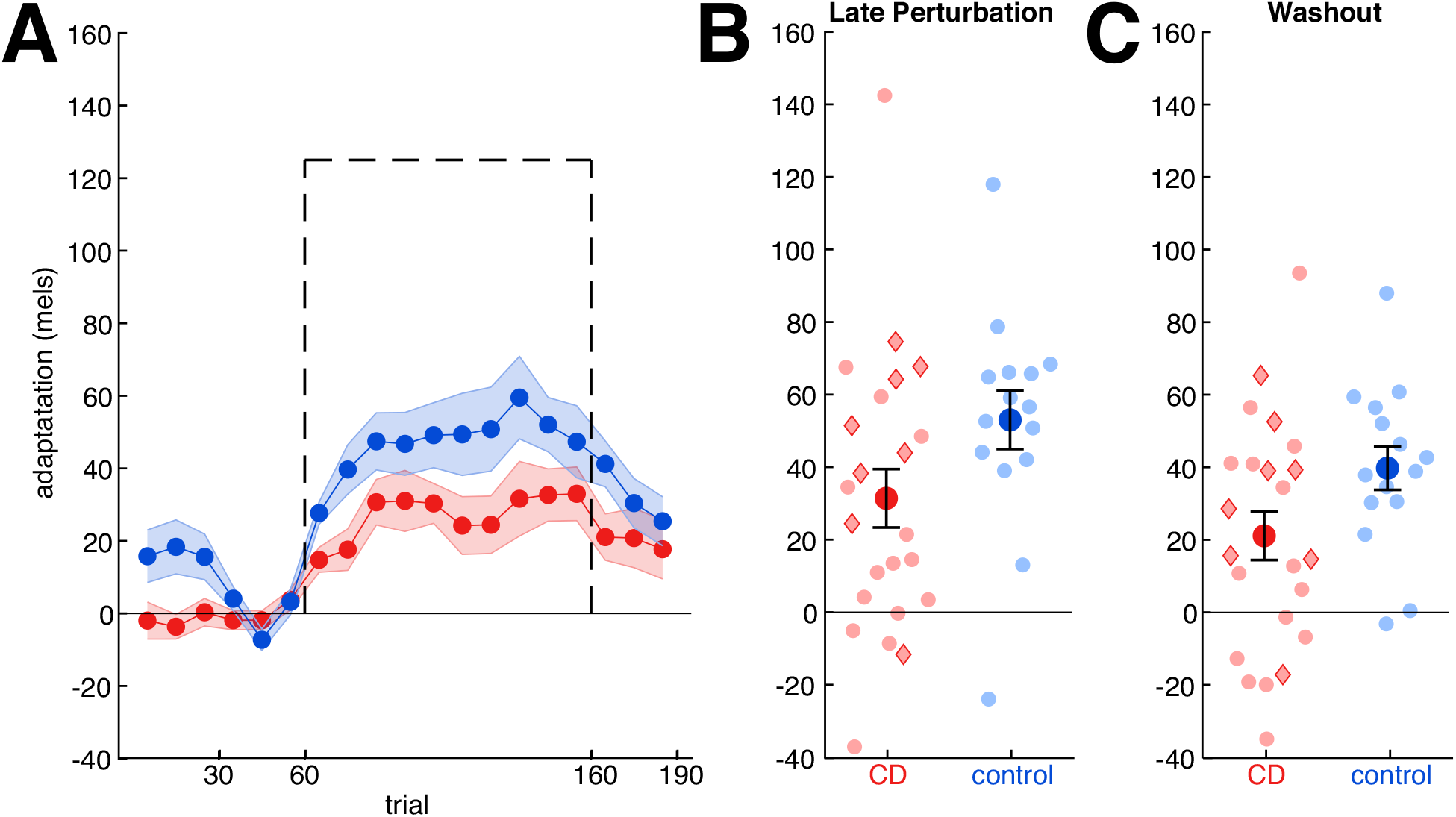
In the speech adaptation task, the CD group adapted less than controls. **A:** Change in first formant over the course of the experimental condition. Means and standard errors are shown for control (blue) and CD participants (red). For each participant, the change was calculated relative to their mean F1 value during the second half of the baseline phase, with positive values corresponding to changes in the opposite direction of perturbation. The perturbation is shown with a dashed black line (sign flipped). **B**,**C:** Group means ± standard error (solid dots) and individual data points (semi-transparent dots) of adaptive response during the last 10 trials of the perturbation phase and first 10 trials of the washout phase. Red diamonds indicate individuals with SCA6 and SCA8; red circles indicate the other CD participants.

We first confirmed that both groups adapted during the perturbation phase (CD: 31 ± 39 mels, *t*(1,22) = 3.9, *p* = 0.0007, *g* = 0.79; control: 53 ± 31 mels, *t*(1,14) = 6.6, *p* < 0.0001, *g* = 1.61; Figure 4B) and maintained their adapted state during the initial part of the washout phase (CD: 21 ± 32 mels, *t*(1,22) = 3.1, *p* = 0.005, *g* = 0.64; control: 40 ± 23 mels, *t*(1,14) = 6.6, *p* < 0.0001, *g* = 1.62; Figure 4C). The CD group exhibited less adaptation than the control participants (*F*(1,36) = 4.7, *p* = 0.036, *η*^*2*^ = 0.12), and this difference was similar in both phases (Group x Phase interaction: *F*(1,36) = 0.06, *p* = 0.80, *η*^*2*^ = 0.002). In sum, we find that sensorimotor adaptation in speech is impaired in individuals with CD, similar to the impairment observed for reach adaptation in Condition 1 and consistent with previous findings (Parrell et al., 2017).

### Condition 4: Feedback control in speech

Online corrective responses for speech were operationalized as the change in F1 during the time window from 200-300 ms after vowel onset in response to an upward or downward perturbation of F1, randomized across trials. The data are plotted relative to F1 values measured on unperturbed trials. As can be seen in Figure 5A, the non-predictable auditory perturbations resulted in compensatory responses that opposed the perturbation in both groups. The magnitude of the corrective response (7.6% in controls, 9.2% in the ataxic group) was similar to that typically observed in response to auditory perturbations of speech and much lower than observed for the non-predictive perturbations during reaching used in Condition 2.

**Figure 5:**
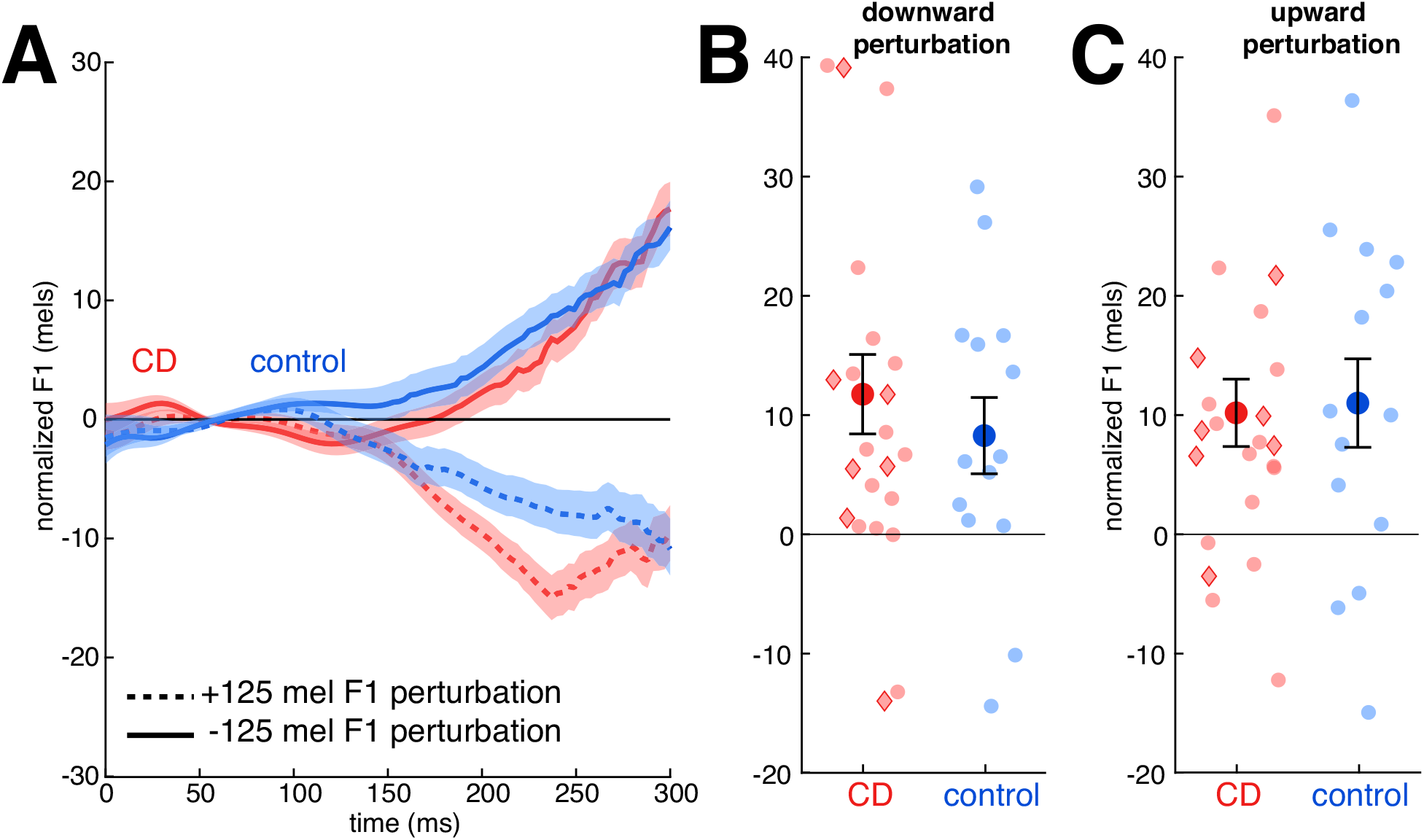
In the speech feedback control task, both the CD group and controls corrected for formant perturbations in both directions, with no difference between the two groups. **A:** Normalized formant trajectories in response to a downward F1 perturbation (solid lines) or upward perturbation (dotted lines) in the control (blue) and CD (red) groups. Shaded area indicates standard error. Data are normalized to the formant trajectories produced in trials with no perturbation. **B, C**: Magnitude of F1 values in trials with a downward (**B**) or upward (**C**) perturbation, averaged over the time window spanning from 150-300 ms following onset of vocalic portion of the utterances. Compensatory responses are expressed as positive values in both cases. Group means ± standard error are shown as solid dots; Semi-transparent dots represent individuals. Red diamonds indicate individuals with SCA6 and SCA8; red circles indicate the other CD participants.

The change on perturbed trials was significantly larger than 0 in response to upward and downward perturbations in the CD group (up: 10.2 ± 13.6 mels, *t*(1,22) = 3.6, *p* = 0.002, *g* = 0.73; down: 11.8 ± 16.0 mels, *t*(1,22) = 3.5, *p* = 0.002, *g* = 0.71) and control group (up: 11.0 ± 14.4 mels, *t*(1,13) = 2.9, *p* = 0.02, *g* = 0.72; down: 8.2 ± 12.4 mels, *t*(1,13) = 2.9, *p* = 0.02, *g* = 0.72) (Figure 5B, C). While the mean values were larger in the CD group and individuals in this group showed the largest compensatory response, the difference between the two groups was not significant (*F*(1,35) = 0.35, *p* = 0.56, *η*^*2*^ = 0.009). There were no differences between the responses to the two perturbation directions (*F*(1,35) = 0.1, p = 0.74, *η*^*2*^ = 0.003) and the Group x Direction interaction was not significant (*F*(1,35) = 0.07, *p* = 0.80, *η*^*2*^ = 0.002). These results suggest feedback gains for auditory perturbations in speech are similar in individuals with CD and healthy controls, consistent with the reaching results in Exp 2. Of note, the null effects here are inconsistent with the results from previous studies involving speech articulation (Parrell et al., 2017) and vocal pitch production (Houde et al., 2019; Li et al., 2019) in which individuals with CD were found to show an enhanced feedback response.

### Feedforward and feedback control within and across motor domains

By testing each participant in all four conditions, we can compare the measures of feedback and feedforward control within and across task domains. Because our focus in this analysis is how deficits in these domains may be correlated in individuals with cerebellar degeneration, we limit this analysis to the 22 individuals with CD.

The between-domain comparisons assess the similarity of impairment (or lack thereof) between reaching and speech. Although the ataxic group adapted less than the controls at the group level, there was no significant correlation within the ataxic group between the magnitude of adaptation between speech and reaching (Figure 6A). Similarly, the magnitude of feedback-based compensation (**Error! Reference source not found**.B) was unrelated across motor domains.

**Figure 6:**
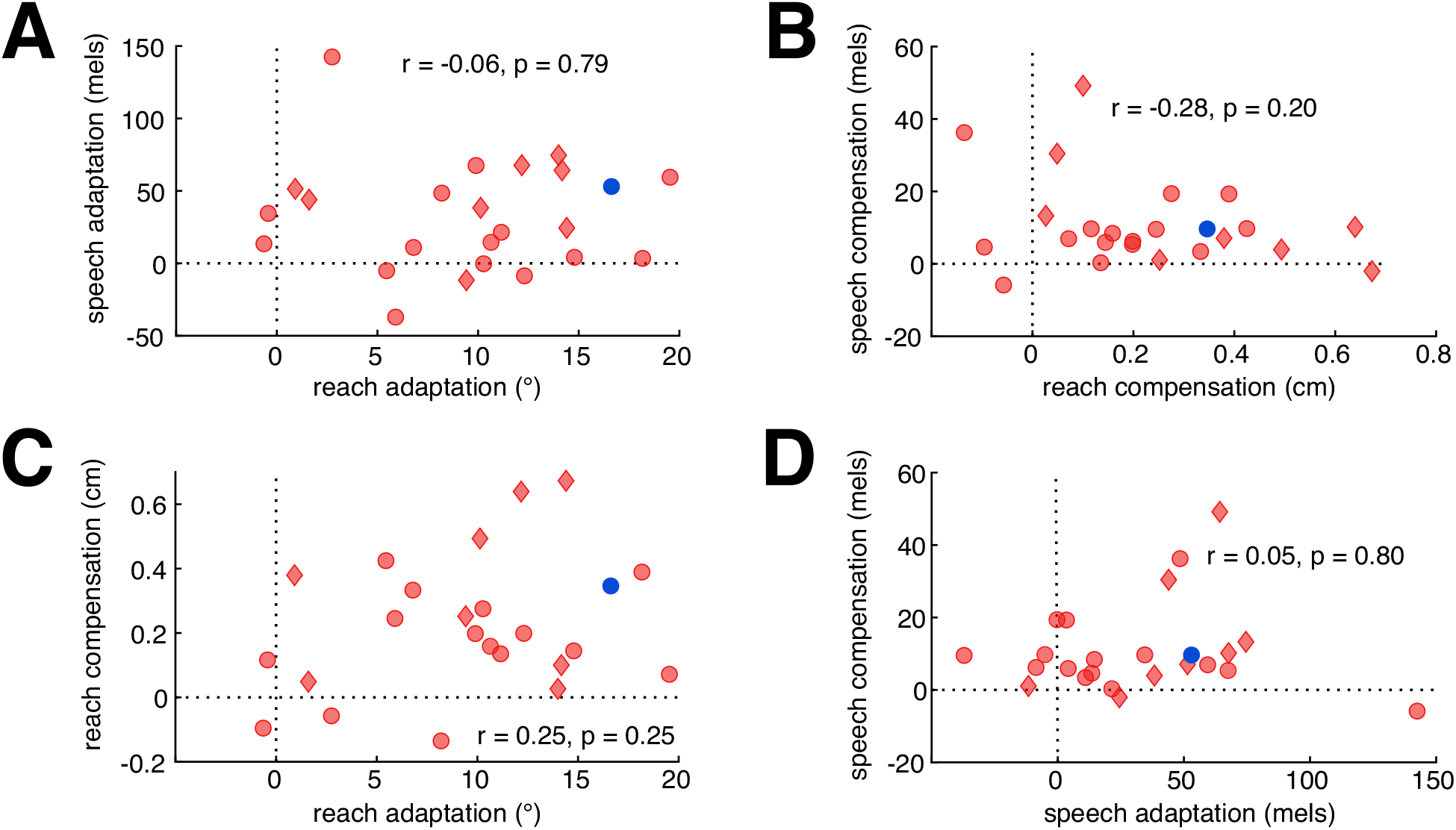
Correlational analyses of feedforward adaptation and feedback-based compensation in individuals with cerebellar degeneration. Within the CD group, there were no correlations in behavioral measures obtained across (**A-B**) or within (**C-D**) motor domain. Plots show individual CD participants as red symbols (red diamonds, SCA6 and SCA8; red circles, other CD participants) and the mean of the control participants as a blue dot. To simplify interpretation, the sign of compensation in reaching has been flipped such that larger positive values reflect greater compensation, as for all other measures. **A:** Adaptation in reaching and speech. **B:** Compensation in reaching and speech. **C:** Adaptation and compensation in reaching. **D**: Adaptation and compensation in speech.

The within domain correlations provide a test of the hypothesis that feedback gains may increase in response to impaired feedforward control: in this case, we would expect a negative correlation between these measures such that individuals who are more impaired in feedforward control should be more likely to exhibit a greater reliance on feedback control. This relationship might hold even if there is no overall increase in compensatory responses when comparing the CD and control groups. Contrary to this prediction, the relationship between feedforward adaptation and feedback-based compensation was not significant in either reaching (**Error! Reference source not found**.C) or speech (**Error! Reference source not found**.D).

We additionally tested whether any of the measures of feedforward and feedback control correlated with ataxia severity as assessed with the SARA. Neither a summary measure of overall upper limb ataxia nor intentional tremor severity correlated with reach adaptation or compensation (all *p* > 0.2). Similarly, overall speech impairment was not correlated with speech compensation (*r* = -0.24, *p* = 0.26). We did observe a positive correlation between the SARA measure of speech impairment and speech adaptation, with lower levels of adaptation to the auditory perturbation associated with greater speech impairment, though this was not significant after correction for multiple comparisons (*r* = 0.45, *p* = 0.03). Participant age was not correlated with either SARA scores in the CD group or the behavioral measures in either the CD group or controls (all *p* > 0.31).

## Discussion

We tested a group of CD participants across a series of tasks to evaluate the impact of cerebellar degeneration on feedforward and feedback control in two motor domains, reaching and speech. Individuals with cerebellar degeneration showed a marked impairment in feedforward control relative to controls, manifest as reduced adaptation in response to a sensory perturbation that remained constant from trial to trial. Feedback control, measured as the on-line response to a variable perturbation, was intact in both task domains.

### Multi-modal impairment in feedforward control in individuals with cerebellar degeneration

Our principle positive result, that individuals with cerebellar degeneration are impaired in adapting their motor behavior in the presence of sensory prediction errors, is consistent with prior neuropsychological studies involving upper limb control and speech. These results are consistent with the hypothesis that the cerebellum provides a domain-general mechanism for generating sensory predictions and using error information to keep this predictive system well-calibrated. We assume this process operates in an automatic and implicit manner. Adaptation in speech is highly likely to be an implicit process (Munhall et al., 2009; Kim and Max, 2020; Lametti et al., 2020) as are adaptive responses in reaching in response to perturbations similar to those employed in the current study (Bond and Taylor, 2015; Morehead et al., 2015; Werner et al., 2015).

While adaptation was impaired in both reaching and speech at the group level, we did not observe correlations between the two motor domains. It is possible that the null results in the correlational analyses reflect statistical limitations with our design. Given that we only tested each condition a single time, we do not have estimates of the reliability of our estimates of adaptation and compensation in each domain, and the upper bound for a correlation analysis is set by the reliability of each measure. There is also concern that we lack sufficient power given our sample size of 22 individuals with CD, although Bayes factor values (0.46-0.88) indicate that we have some evidence against the predicted correlations. However, previous work has identified a similar dissociation in individuals with CD between adaptation to dynamic (force-field) and kinematic (visuomotor rotation) perturbations during reaching in both behavior and cerebellar localization (Rabe et al., 2009; Donchin et al., 2012). Our results provide further evidence that dissociable regions of the cerebellum are involved in different forms of adaptation. Given existing evidence of speech localization in the cerebellum (Urban et al., 2003; Ackermann, 2008; Peeva et al., 2010; Mariën et al., 2014), we would anticipate that deficits in speech adaptation would be associated with intermediate portions of lobule VI bilaterally, relative to more lateralized regions of lobule V/VI that have been associated with deficits in limb adaptation (Rabe et al., 2009; Donchin et al., 2012).

Future work examining patterns of cerebellar damage in patients will be needed to directly test this hypothesis. A detailed analysis of structural MRIs would also provide the means to assess the contribution of extracerebellar structures to feedforward and feedback control. Given our heterogenous sample, we expect that the pathology in at least some of the participants with CD extended to extracerebellar structures. While this qualifies our assumption that the observed deficits are associated with cerebellar dysfunction, we did observe that individuals with genetic variants in which the pathology is relatively restricted to the cerebellum (SCA6 and SCA8) performed qualitatively similarly to the rest of the sample. As such, we think it reasonable to attribute the impairments in feedforward control to the cerebellar pathology.

### Intact, but not enhanced feedback control in individuals with cerebellar degeneration

Feedback-based corrections for errors were similar in magnitude between individuals with cerebellar degeneration and controls for both speech and reaching. Although this result agrees with estimates of feedback gains in a continuous visuomotor tracking task (Zimmet et al., 2020), the absence of a difference between the CD and control groups on the speech task fails to replicate our previous finding showing an enhanced feedback response (Parrell et al., 2017). Our previous results had led to the hypothesis that the enhanced feedback response reflected a compensatory mechanism to help offset the disruptive effects of impairment in feedforward control. The failure to observe enhanced feedback in the current study in both domains argues against this compensatory hypothesis. The absence of a correlation between the feedforward and feedback measures within both motor domains also argues against a compensatory hypothesis.

There are a few issues to consider in terms of the discrepancy between our results and previous studies as well as the interpretation of the null results regarding the compensatory hypothesis. First, we may have failed to find any changes in feedback control in individuals with CD simply due to sampling issues. Our sample of this population (n = 22) is consistent with, or larger than, many previous studies (Morton and Bastian, 2006; Parrell et al., 2017; e.g., Houde et al., 2019; Li et al., 2019; Zimmet et al., 2020), and we were adequately powered to detect relatively large between-group effects (.75 power to detect *d* of .8). However, the effect size of the expected increase in feedback gains in certain domains, as in our previous work on oral articulation, may be relatively small. Although we did find a numerically larger feedback response for speech in the CD group compared to controls, this difference was smaller than in our previous work (Parrell et al., 2017). Thus, we may simply have been under-powered to detect changes associated with CD.

Second, it may be that increased feedback gains are a secondary, and somewhat sporadic, effect of cerebellar degeneration. The consistent and striking deficit in feedforward control points to inaccurate or attenuated predictive signals. In this case, some individuals may learn to rely more on sensory feedback to help with movement accuracy as a compensatory mechanism for impairments in feedforward control. It may be that our sample did not include enough participants with altered feedback gains to show an effect at the group level. If this is the case, we might still expect to find some evidence that individuals with larger feedback gains have greater impairments in feedforward control, even if there were no group differences. However, we found no evidence of this negative correlation, making this account less likely.

A final potential explanation is that increases in feedback gains are domain-specific while impairments in feedforward control are more general, at least at the group level. The strongest example of higher feedback gains in individuals with CD comes from work on pitch control, where the on-line response to a pitch perturbation is roughly twice as large in a CD group compared to a control group (Houde et al., 2019; Li et al., 2019). This increase may result from an inherent reliance on auditory feedback for pitch control, compared to higher reliance on feedforward control for speech and reaching, as evidenced by the fact that pitch control rapidly degenerates after post-lingual hearing loss while oral articulation is better maintained (Cowie and Douglas-Cowie, 1983; Lane and Webster, 1991). Thus, cerebellar degeneration may cause increased feedback gains only in those motor domains which are already primarily reliant on sensory-feedback-based control.

These results raise a broader question regarding the role of the cerebellum in feedback control. Traditionally, the cerebellum has been thought to be involved in both feedforward and feedback aspects of motor control (e.g., Herrick, 1924; Guenther, 2016). However, our results suggest limited changes in feedback control in individuals with CD. It is possible that feedback control and feedforward control are localized in different cerebellar subregions (Evarts and Thach, 1969; Allen and Tsukahara, 1974), and that cerebellar degeneration in our sample was localized to areas critical for feedforward control. Alternatively, it may be that the cerebellum plays a central role in feedforward control but a more modulatory role in feedback control.

Specifically, the cerebellum may be needed to generate appropriate corrective commands in a dynamic context, when the state of the moving body must be estimated; in contrast, it may not be essential under static conditions when the state can be more directly inferred from sensory feedback (Wolpert et al., 1998; Miall et al., 2007). Consistent with this view, individuals with CD produce appropriately-scaled changes in precision grip force to discretely-presented changes in load but not to continuously varying, but unpredictable, load changes (Nowak et al., 2004; Brandauer et al., 2010). Future work could test these possibilities using MRI imaging coupled with more complex reaching and speech tasks.

### Conclusions

Adaptation of feedforward control based on sensory errors is impaired in individuals with cerebellar degeneration in both reach and speech. Contrary to our initial hypothesis and data from vocal pitch control, we found no evidence for increased feedback gains in either domain. However, these results, together with those from a recent study of upper extremity control (Zimmet et al., 2020), motivate further investigation into how feedback gains may be differentially affected by the specific demands of different motor tasks, as well as to determine the variability in feedback control associated with cerebellar dysfunction.

## Acknowledgments

This work was funded by grant R01 DC017091 from the NIDCD to BP and RI, R35 NS116883 from the NINDS to RI, K12 HD055931 from the NICHD to HEK, and a core grant U54 HD090256 from the NICHD to the Waisman Center.

